# Lifespan Analysis of the Lung Epithelium Reveals Inflammatory Reprogramming and Regenerative Decline

**DOI:** 10.64898/2026.06.01.729288

**Authors:** Gowtham Boosarpu, Eva M. Guenther, Carina Steinchen, Maria Camila Melo Narvaez, Camille Barro, Turgud Nazarli, Antonia Müller, Francesco Campi, Friso F. Bratzel, Laura-Marie Twardowski, Qianjiang Hu, Efthymios Fousekis-Papakonstantinou, Roxana M. Wasnick, Hosam Shams-Eldin, Anne Hilgendorff, Anna Lena Jung, Ali Önder Yildirim, Marie Piraud, Bernd Schmeck, Melanie Königshoff, Tobias Stöger, Miguel A. Alejandre Alcazar, Carola Voss, Mareike Lehmann

**Affiliations:** Institute for Lung Research, Philipps-University Marburg, German Center for Lung Research (DZL), Marburg, Germany; Institute of Lung Health and Immunity (LHI), Helmholtz Munich, Comprehensive Pneumology Center (CPC-M), German Center for Lung Research (DZL), Neuherberg, Germany; Translational Immunology, School of Medicine and Health, Technical University of Munich (TUM), Munich, Germany; Helmholtz AI, Computational Health Center (CHC), Helmholtz Munich, Neuherberg, Germany; University of Cologne, Faculty of Medicine and University Hospital Cologne, Translational Experimental Pediatrics, Department of Pediatric and Adolescent Medicine, Cologne, Germany; University of Cologne, Faculty of Medicine and University Hospital Cologne, Cologne Excellence Cluster for Aging and Aging-Associated Diseases (CECAD), and Center for Molecular Medicine Cologne (CMMC), Cologne, Germany; Division of Pulmonary, Allergy & Critical Care Medicine, University of Pittsburgh Medical Center, Pittsburgh, PA, United States of America; Animal Experimental Facility, Biomedical Research Center (BMFZ) Philipps-University Marburg, Marburg, Germany; Core Facility Flow Cytometry – Bacterial Vesicles, Philipps-University Marburg, Marburg, Germany; Institute of Experimental Pneumology, University Hospital Munich, Ludwig-Maximilians University, Munich, Germany, Munich, Germany; Centre for Synthetic Microbiology (Synmikro), Philipps-University Marburg, Marburg, Germany; Department of Respiratory, Intensive Care, and Sleep Medicine, Clinic for Respiratory Infections, Philipps-University Marburg, Marburg, Germany; Institute for Lung Health (ILH), German Center for Lung Research (DZL), and Cardiopulmonary Institute (CPI), Justus-Liebig University Giessen, Germany; Hannover Medical School, Clinic for Cardiac, Thoracic-, Transplantation and Vascular Surgery, Leibniz Research Labs for Biotechnology and Artificial Organs (LEBAO), Biomedical Research in End-stage and Obstructive Lung Disease (BREATH), German Center for Lung Research (DZL), Hannover, Germany

## Abstract

Aging is a major risk factor for chronic lung diseases, associated with chronic low-grade inflammation (inflammaging) and impaired epithelial regeneration. How epithelial-intrinsic aging intersects with inflammaging across the lifespan remains poorly understood. Here, we systematically analyzed lung epithelial cells from neonatal, young adult, and aged mice to define age-dependent changes in regenerative capacity.

RNA sequencing revealed lifespan-associated shifts characterized by early repression of developmental and WNT/β-catenin programs and progressive activation of DNA damage, inflammation, and senescence signatures. Functionally, neonatal epithelial cells exhibited markedly enhanced organoid-forming capacity compared with young and aged cells. Aged organoids maintained a pro-inflammatory secretory profile indicative of cell-intrinsic inflammaging, and transfer of the aged secretome or TNF-α to young cultures significantly impaired regeneration. Comparison of freshly isolated cells and long-term organoid cultures revealed sustained repression of regenerative pathways with age, consistent with stable epigenetic imprinting. Pharmacological inhibition of DNA methylation and WNT signalling partially restored regenerative capacity in adult organoids.

Together, these findings identify epigenetic reprogramming and epithelial-intrinsic inflammaging as key determinants of age-dependent regenerative decline.

## Introduction

Aging is a major risk factor for respiratory diseases, with impaired regenerative capacity contributing to the development of chronic lung conditions such as chronic obstructive pulmonary disease (COPD) and idiopathic pulmonary fibrosis (IPF), which together rank among the leading causes of mortality worldwide (Ferrari *et al*, 2024; Gillissen *et al*, 2023). The decline in lung repair mechanisms with age has emerged as a central determinant of disease susceptibility and progression, yet the underlying molecular pathways remain incompletely understood.

Recent work has highlighted the role of aging in limiting regenerative responses in the lung, implicating multiple hallmarks of aging, including stem cell exhaustion, epigenetic reprogramming, and cellular senescence, as well as extrinsic factors such as chronic low-grade inflammation, often referred to as inflammaging (Melo-Narváez *et al*, 2020; Schuliga *et al*, 2021; Han *et al*, 2023; Maté *et al*, 2021; Baker *et al*, 2026). These processes collectively alter the functional capacity of resident progenitor cells and reshape the tissue microenvironment. However, it remains unclear which signalling pathways causally drive the loss of regenerative potential and how these mechanisms are established and maintained over time.

Alveolar epithelial type II (AT2) cells serve as the principal stem cell population of the distal lung, maintaining tissue homeostasis by self-renewal and differentiation into alveolar epithelial type I (AT1) cells following injury (Nabhan *et al*, 2023). These processes are tightly regulated by different cellular signalling pathways, including the WNT/β-Catenin pathway (Bush, 2021). Notably, recent single-cell analyses have identified AT2 cells as one of the most aging-affected cell populations in the human lung (Man *et al*, 2026). Functional studies further demonstrate a progressive decline in AT2 regenerative capacity with age, as evidenced by reduced proliferative responses following injury, diminished organoid-forming efficiency, and impaired alveolar regeneration in vivo (Wang *et al*, 2023; Lehmann *et al*, 2020; Yazicioglu *et al*, 2020; Rowbotham *et al*, 2023; Zhuang *et al*, 2025). In contrast, the younger lung exhibits a remarkable capacity for repair, with neonatal epithelial cells displaying high plasticity and regenerative potential (Penkala *et al*, 2021). Despite this, the molecular mechanisms that govern epithelial regeneration early in life, and how these processes are altered during aging, remain poorly defined.

Emerging evidence suggests that early-life exposures, including genetic predisposition, maternal smoking, mechanical ventilation, and respiratory infections, can induce long-lasting alterations in lung structure and function, thereby predisposing individuals to chronic lung disease later in life (Bush, 2021; Hopkinson *et al*, 2024; Kirkeleit *et al*, 2023; Mocelin *et al*, 2022; Duan *et al*, 2021; Griese *et al*, 2024; Saeed *et al*, 2023). These observations highlight the importance of understanding how epithelial cell regeneration progressively erodes with aging.

Here, we use primary cells and a murine lung organoid model based on neonatal, young and old epithelial cells to investigate how inflammation, epigenetics and aging collectively converge to alter regeneration of lung epithelial cells over the lifespan, and to identify mechanisms that may be targeted to restore regenerative potential in the aging lung.

## Methods

### Animals

Female C57BL/6J mice of the following ages were used in the experiments: between 8 and 20 weeks old for young mice and between 66 and 93 weeks old for old mice, and for neonatal mice, pups of undetermined sex between 5 and 14 days old were used. Mice were housed in individually ventilated cages under specific pathogen free conditions. Three to four animals were housed in one cage (Tecniplast Greenline GM 500, 501 cm² floor space) with a 12/12 – hour light/dark cycle. The barrier animal rooms were fully air-conditioned, with 20 – 24 °C temperature and 45 – 65 % humidity. Animals had ad libitum access to sterile filtered water and a standard diet for rodents (Altromin 1314). To improve housing conditions, the cages were equipped with laboratory animal bedding (wood fiber/wood chips; Lignocel Select Fine, SAFE), autoclaved nesting material (Arbocel Crinclets, SAFE), plastic mouse houses and wooden bricks.

Hygiene monitoring was carried out at least four times a year in accordance with the current FELASA recommendation. In the animal housing areas equipped with IVC systems, exhaust dust from the IVC ventilation units was tested for all FELASA-listed pathogens using PCR. In addition, animals with clinical abnormalities were examined using classical examination methods (bacteriology, parasitology, pathology). During the time of these experiments, all health monitoring reports were negative and did not affect the results of this study.

### Lung epithelial cell isolation

Mice were anaesthetized with 100 µl of a lethal anesthesia mixture of ketamine (150 mg/kg) and xylazine (10 mg/kg) and killed painlessly by blood withdrawal at the inferior vena cava. Blood was flushed from the lungs with PBS, and the lungs were inflated intratracheally with Dispase (50U/ml, Corning) followed by 1 % low gelling temperature agarose (Sigma-Aldrich), excised and incubated in Dispase (50U/ml, Corning) for 40 minutes at RT. For neonatal mice, lungs were incubated in Dispase (50U/ml, Corning) without intratracheal instillation. The lungs were then minced and filtered through 100 and 40 µm nylon meshes. Red blood cells were depleted with RBC lysis buffer (Invitrogen). Using magnetic bead sorting, macrophages and white blood cells as well as endothelial cells were removed using CD45 and CD31 beads (Miltenyi Biotec), respectively. Epithelial cells were enriched by using CD326 (EpCAM) beads (Miltenyi Biotec) according to the manufacturer’s instructions.

### Organoid culture

For organoid culture, proliferation of MLg (ATCC CCL-206) mouse lung fibroblasts was inactivated with 10 µg/ml mitomycin C (Sigma-Aldrich) for 2 h followed by a 1 h recovery in culture media. 25µl of media containing 10,000 EpCAM^+^ cells was mixed in a 1:1 ratio with growth-arrested MLg CCL206 cells, followed by a mixing with 25µl of growth factor-reduced Matrigel, and seeded into a 96-well plate or an 8-well chamber. The gel was allowed to polymerize at 37°C for 30min. Cultures were maintained in 100 µl DMEM/F12 containing 100 U/ml penicillin and 100 µg/ml streptomycin, 2 mM L-alanyl-l-glutamine (Gibco), Amphotericin B (Gibco), insulin-transferrin-selenium (Gibco), 0.025 µg/ml recombinant human EGF (Sigma Aldrich), 0.1 µg/ml Cholera toxin (Sigma Aldrich), 30 µg/ml bovine pituitary extract (Sigma Aldrich), and 0.01 µM freshly added all-trans retinoic acid (Sigma Aldrich). 10 µM Y-27632 (Tocris), a Rho-associated kinase (ROCK) inhibitor, was added for the first 48 h of culture. Treatments were added to the media from day 0 and with every change of culture media. Treatments include 10 ng/ml TGFβ (R&D Systems), 10 ng/ml IL1β (R&D Systems), 0.05 and 0.5mM Azacytidine (Sigma Aldrich) and 10 mM Lithium Chloride (LiCl) (Sigma Aldrich). Media was changed every 2-3 days.

### High-content imaging of Organoids and Napari organoid counter

Imaging of organoid cultures was performed using the Operetta CLS high-content analysis system (Operetta CLS, PerkinElmer) with Harmony software (PerkinElmer) or the Leica Thunder Imager on day 14 of culture. Quantification, classification into 2 classes (grape-like and cystic-like) and size measurements of organoids were performed using the “Napari organoid counter v0.2.6” after visualizing multi-plane confocal 3D images as Extended Depth of Focus projections in Napari image viewer (Python, <u>HelmholtzAI-Consultants-Munich/napari-organoid-counter: v0.2.6,</u> https://zenodo.org/records/19824587). The Organoid Numbers from all technical and biological replicates were then used to calculate Colony forming efficiency for each group. The same procedure was used for the organoids classified into the grape-like and cystic-like classes.

### Immunofluorescence

All antibodies used for Immunofluorescence are listed in Supplementary Table 1

### Immunofluorescence of mouse lung organoids

Organoids were cultured in 8-well chamber slides for immunofluorescence analysis (Nunc Lab-Tek Chamber Slide System, 8-well, Permanox slide, 0.8 cm^2^ /well). After 14 days of culture and treatments, organoids were fixed with ice cold methanol and acetone (1:1v/v) for 5 minutes at – 20°C. After washing with PBS, organoids were stained with the respective primary antibodies in buffer containing 0.1% BSA and 0.1% Triton X-100 overnight at 4°C. Cells were washed 3 times with PBS and incubated with the respective fluorescent conjugated secondary antibody at a dilution of 1:500 and DAPI diluted 1:1,000 the next day overnight at 4°C. On the third day, organoids were washed with PBS again. The growth chamber was removed, and finally each well was mounted with fluorescence mounting media (Dako) and covered with a coverslip. Slides were stored at 4°C until imaging. Imaging was performed using a confocal laser scanning microscope (CLSM) Zeiss LSM 880 with Airy scan and edited afterwards using ZEN 2.5 software (Zeiss).

### Embedding and staining of murine lung tissue for structural lung cell markers

Lung tissue samples were washed once with 1X PBS, fixed in 4% PFA in PBS at 4°C overnight and kept in PBS until used. Fixed lung tissues were transferred to embedding cassettes and processed by a Microm STP 420D Tissue Processor (Thermo Scientific, USA) following next steps: 50% EtOH (1 cycle, 60 min), 70% EtOH (1 cycle, 60 min), 96% EtOH (2 cycles, 60 min each), 100% EtOH (2 cycles, 60 min each), Paraffin (1 cycle, 30 min), and Paraffin (3 cycles, 45 min each). Then, tissue was embedded in paraffin Type 3 (Thermo Scientific, USA) using the Modular Tissue Embedding Center EC 350 (Thermo Scientific, USA). Formalin-fixed, paraffin-embedded (FFPE) sections were cut at 3μm using a Hyrax M55 microtome (Zeiss, Germany) mounted on slides, dried overnight at 40°C. Subsequent steps differed across two protocols: To stain for structural lung cell markers FFPE sections were deparaffinized, rehydrated and exposed to heat-induced antigen retrieval (10 mM citrate buffer, pH = 6.0, 1 cycle at 125°C for 30 s, 1 cycle at 90°C for 10 s). Slides were washed twice with 1X PBS and blocked for 1 h at RT in a humid chamber with 10% Normal Donkey Serum in DAKO Antibody Diluent (Agilent Technologies, USA). Primary antibodies were diluted in DAKO Antibody Diluent (Agilent Technologies, USA) added to the samples and incubated at 4°C, overnight in a humid chamber. The next day, samples were washed twice in 1X PBS for 5 min and incubated with secondary antibodies (1:250) and DAPI (1:400) prepared in 1% BSA in PBS and incubated for 2-h at RT in a humid chamber. Finally, samples were mounted using DAKO Fluorescence mounting medium (Agilent Technologies, USA) and imaged using an AxioImager (Zeiss) at 20X magnification. Qupath was used for automatic quantification of the percentage of positive cells based on cell count from nuclear DAPI signal. Three random regions of interest from four biological replicates were analyzed for the quantifications. To stain for nuclear markers of DNA damage and repair, FFPE sections were deparaffinized in Neoclear (Sigma-Aldrich) for three changes of 15 minutes each and rehydrated in a descending ethanol series (100 %, 96 %, 80 %, and 70 %) for 1 minute each and washed in distilled water for 1 minute. Sections were permeabilized with 1 % Triton X-100 in PBS for 10 minutes at RT beforehand. Autofluorescence was reduced by incubation with Reagent A from the MaxBlock Autofluorescence Reducing Kit (MaxVision Biosciences) for 5 minutes at RT. Slides (Epredia, #10149870) were washed in 60% ethanol for 1 minute, distilled water for 5 minutes, and three times in PBS. Sections were permeabilized with 1 % Triton X-100 in PBS for 10 minutes at RT. Antigen retrieval was performed by heating the slides to 90–120 °C in 10 mM citrate buffer (pH 6; Dako) in a water bath (Braun) for 25 minutes, followed by cooling for 20 minutes at RT. Endogenous peroxidase activity was blocked by incubation with 3 % peroxidase (H_2_O_2_)) in distilled water for 10 minutes at RT. Sections were incubated with SEA BLOCK (Thermo Fisher Scientific) for 60 minutes at RT. Primary antibody incubation was performed overnight at 4 °C in a humidified chamber using the following antibodies diluted in antibody diluent (Dako). After washing, sections were incubated for 60 minutes at RT with the secondary antibody. Sections were washed in PBS and incubated with DAPI (Sigma-Aldrich, 1:1000) diluted in Reagent B from the MaxBlock kit (MaxVision Biosciences) for 10 minutes at RT. Slides were washed three times in distilled water, mounted with glass coverslips using Fluoromount (Sigma-Aldrich) and imaged using an Olympus IX81 at 40X magnification. Quantification was performed by manually counting marker-positive (+) cells relative to all DAPI (+) cells and either SFTPC (+) or ABCA3 (+) cells (%) on ImageJ. Three random regions of interest from four biological replicates were analyzed for the quantifications.

### RNA sequencing of EpCAM^+^ cells

Cell lysis from isolated EpCAM^+^ cells was done using RLT Plus Lysis Buffer (Qiagen). RNA isolation was performed using the RNeasy Mini Kit (Qiagen) according to the manufacturer’s instructions and eluted in 30 µl water. RNA concentration was determined using a spectrophotometer (DeNovix, USA) or NanoDrop 1000, and samples were stored at –80 °C until sequencing. Samples were sent to the Helmholtz Core Facility Genomics (Germany). The mRNA was enriched with oligo(dT) magnetic beads, and the libraries were prepared with the Illumina® Stranded mRNA Prep, Ligation Kit. Samples were ligated with adapters and sequenced using paired-end strand-specific sequencing with the NovaSeq6000 S2 Flowcell and 200 cycles (Read1: 100-Index i7: 10 Index i5: 10 Read2: 100, 30 million reads/sample). Adapters were trimed using Cutadapt (v.2.10) using the adapters sequences: –A CTGTCTCTTATA and –a CTGTCTCTTATA. Low quality reads were removed with cut off –q 15. Then, reads were aligned against GRCm39 (mouse) using HISAT2 (v2.2.1). Counts were obtained from the mapped reads using featureCounts (subread, v2.0.1) using the parameter “-stranded reverse”. Gene annotation was added according to Ensembl version 108. Downstream analyses were conducted in R (v4.5.2) and RStudio (v2026.01.1 Build 403). Raw counts from neonates, young and old were merged and corrected for batch bias using ComBatseq and surrogate variables (svseq) functions in sva package (v3.58.0). Then, differential expression analysis was done using the DESeq2 package (pAdjustMethod = “BH”, alpha = 0.05, v.1.50.2) and shrinkage of the Log-fold change (LFC, ashr method). Differentially expressed genes (adjusted p-value< 0.05, LFC>1) were extracted and used for downstream analysis and data exploration. Gene set enrichment analyses were performed using fsgea (v1.36.0), DOSE (v4.4.2), and Cluster Profiler (v4.18.4) packages.

### RNA sequencing of Organoids

Organoids were harvested and lysed by pooling 4 wells in 1ml of Trizol (Qiagen). Then, one 5 mm stainless steel bead (Qiagen, USA) was added to each sample, and tissue was homogenized using a TissueLyser II (Qiagen, USA) 3 times at 30 Hz for 30 seconds. The samples were then incubated at RT for 5 mins, following which chloroform was added. After mixing by vortexing for 15 seconds and another incubation of 5 mins at RT, the samples were then centrifuged at 4,000 x g/RCF at RT. gDNA was then eliminated by transferring the colorless upper phase to a gDNA eliminator column. (Qiagen, USA). The flow through was then mixed with an equal volume of 100% EtOH and loaded onto a RNeasy Spin column (Qiagen, USA) and centrifuged at more than 8000 x g/RCF for 15 seconds at RT. The column was then sequentially washed with RW1 buffer (16000 x g/RCF for 15 seconds), RPE buffer(16000 x g/RCF for 15 seconds) and RPE buffer again (16000 x g/RCF for 2 minutes). RNA was then eluted in 30 µl water. RNA concentration was determined using a NanoDrop 1000, and samples were stored at –80 °C until sequencing. RNA quantity and quality were assessed by Bioanalyzer, with no significant differences observed across conditions. The mRNA was enriched from total RNA using poly-T oligo-attached magnetic beads and fragmented prior to cDNA synthesis. First-strand cDNA was synthesized using random hexamer primers, and second-strand cDNA was synthesized incorporating dUTP in place of dTTP. Following end repair, A-tailing, adapter ligation, size selection, USER enzyme digestion, amplification, and purification the directional library was ready, which was then validated using Qubit and real-time PCR for quantification and bioanalyzer for size distribution detection. Validated libraries were then pooled, and sequencing was performed on the Illumina NovaSeq X Plus Series (PE150) platform according to effective library concentration and target data output. Ensembl version 108 was used for gene annotation to enable quantification. Downstream analyses were conducted in R (v4.5.2) and RStudio (v2026.01.1 Build 403). Raw counts from young and old organoids were merged and corrected for batch bias using ComBatseq and surrogate variables (svseq) functions in sva package (v3.58.0). Then, differential expression analysis was done using the DESeq2 package (pAdjustMethod = “BH”, alpha = 0.05, v.1.50.2) and shrinkage of the Log-fold change (LFC, apeglm method). Differentially expressed genes (adjusted p-value< 0.05, LFC>1) were extracted and used for downstream analysis and data exploration. Gene set enrichment analyses were performed using fsgea (v1.36.0), DOSE (v4.4.2), and Cluster Profiler (v4.18.4) packages.

### Quantitative real-time PCR

Lysis of organoids was performed with peqGOLD TriFast (VWR Life Science) and RNA isolation was performed using the RNeasy Mini Kit (Qiagen) according to the manufacturer’s instructions. Transcription of RNA to cDNA was done by reverse transcriptase using the High-Capacity cDNA Reverse Transcription Kit (Thermo Fisher Scientific). cDNA was then used to analyze the target gene expression by quantitative real-time PCR with SYBR Green (Thermo Fisher Scientific).

All primer sequences used for qPCR are listed in Supplementary Table 2

### Multiplex cytokine assay

Organoid supernatants were collected at day 13 or 14 and stored at –20 °C. Concentrations of 13 inflammatory cytokines, namely IL-1α, IL-1β, IL-6, IL-10, IL-12p70, IL-17A, IL-23, IL-27, MCP-1, IFN-β, IFN-γ, TNF-α, and GM-CSF, were assessed using the LEGENDplex Mouse Inflammation Panel 13-plex with V-bottom Plate (BioLegend, Cat No: 740446) according to the manufacturer’s instructions. Flow cytometry analysis was performed on a BD FACSymphony™ A1 (BD Biosciences). Recorded data was analyzed using the LEGENDplex Data Analysis Software (BioLegend).

### Statistical Analysis

All bar graphs and heatmaps were statistically analysed using GraphPad Prism v10. Normality was tested using the Shapiro-Wilk test. Quantitative data is represented as mean ± sd. Group comparisons were analyzed using a one sample *t*-test or one-way analysis of variance (ANOVA), as appropriate. All statistical methods and significance values are indicated in the figure of the respective legends. p<0.05 was considered statistically significant.

## RESULTS

### AT2 cells exhibit markers of DNA damage and senescence in aged mice

The distal lung epithelium is a hotspot of injury linked to different CLDs (Baker *et al*, 2026). To determine the occurrence of aging-associated hallmarks in this region across the lifespan, we performed immunofluorescence staining of mouse lungs at neonatal, young adult and old stages. Structural alveolar epithelial cell markers including podoplanin (Pdpn) for type 1 alveolar epithelial cells (AT1) as well as, lysosome-associated membrane glycoprotein 3 (Lamp3) and pro-surfactant protein C (pro-SPC) for alveolar epithelial type 2 cells (AT2) increased from neonates to adult mice but no significant differences were detected between young and old aged mouse lungs (Supplementary Figure 1a-d). Notably, keratin 8 (Krt8) in the alveolar regions, marking alveolar differentiation intermediate (ADI) cells transitioning from AT2 to AT1 (Strunz *et al*, 2020) showed an age-associated increase (Fig 1a, b), suggesting age-related changes in epithelial differentiation. Senescence marker p21 showed a trend toward increase from neonatal to young and old mice (Fig 1c, d). In parallel, markers of genomic instability including DNA-RNA-Hybrids (R-Loop) as well as DNA damage signalling (as marked by phosphoATM, γH2Ax) increased over age and were predominantly detected in AT2 epithelial cells (Fig 1e-j).

**Figure 1:**
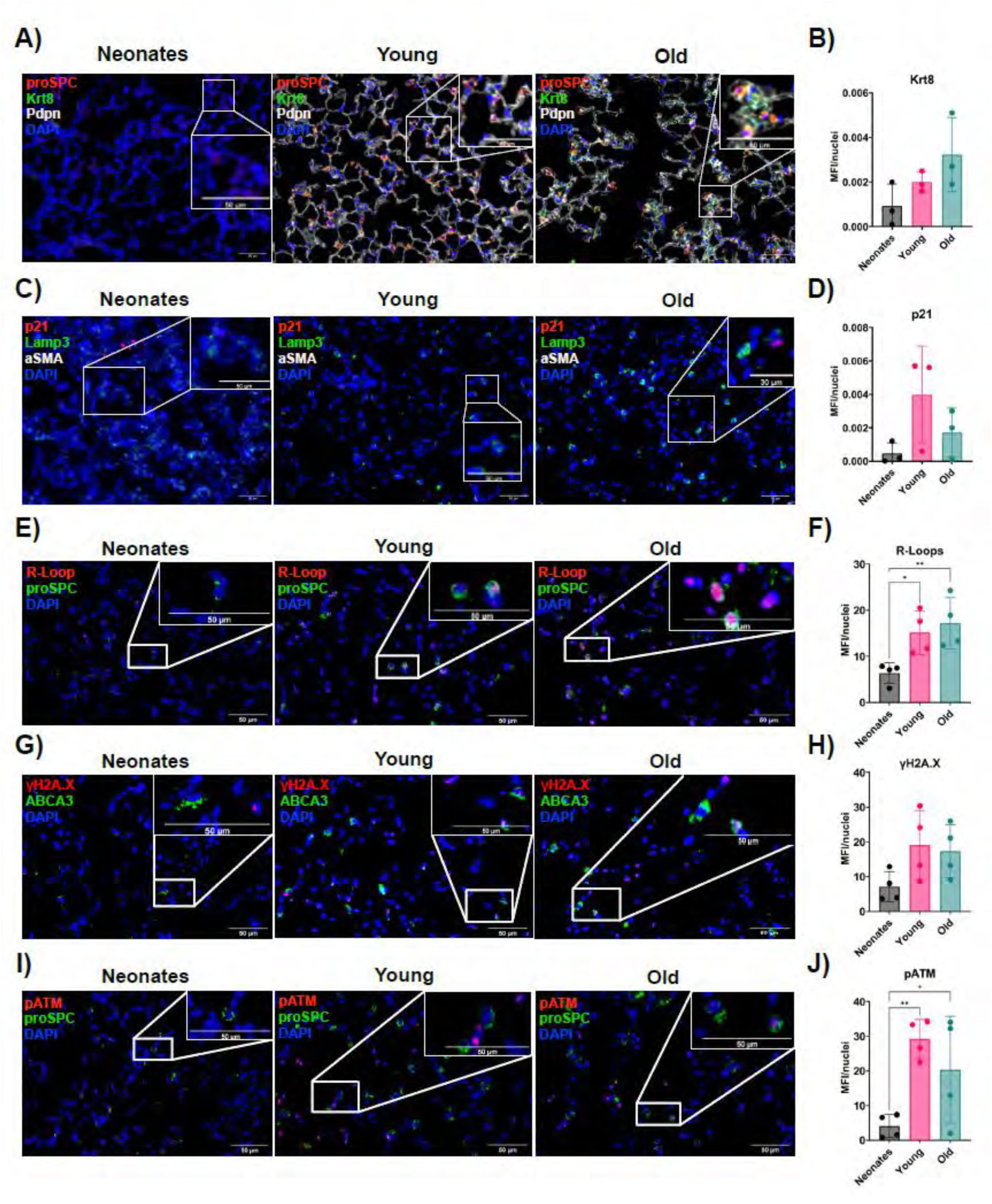
Age-associated accumulation of DNA damage and senescence markers in lung epithelium: **A-K**) Representative images and quantification of mean fluorescence intensity (MFI) normalized to nuclei of immunofluorescence staining of lungs from neonatal, young and old mice stained for **A, B)** DAPI (blue), Krt8 (green), pro-SPC (red) and Pdpn (white), **C, D)** DAPI (blue), Lamp3 (green), p21 (red) and aSMA (white), **E, F)** DAPI (blue), pro-SPC (green), R-Loops (red), **G, H)** DAPI (blue), pro-SPC (green), γH2A.X (red), **I, J)** DAPI (blue), pro-SPC (green), p-ATM (red). All the images and insets have a scale bar of 50 µm. Single points in bar graphs represent different biological replicates. MFI Data are presented as mean ± s.d. Statistical tests: **B, D)** Ordinary one-way ANOVA followed by Tukey’s multiple comparison test, no significance detected. **F, H, J)** Ordinary one-way ANOVA followed by Sidak’s multiple comparison test (selective), *\* p-value < 0.05, ** p-value < 0.01, *** p-value < 0.001, **** p-value <0.0001*.

### Lung epithelial cells show early transcriptional reprogramming over the lifespan

To determine transcriptional changes in lung epithelial cells across the lifespan, we isolated EpCAM^+^ epithelial cells from lungs of neonatal (n=4, postnatal day 5-7), young adult (n=4, 4 months) and old mice (n=4, 21 months) and performed bulk RNA sequencing. A principal component analysis (PCA) showed clear age-related clustering, mainly separating neonatal from young adult and old cells (Fig. 2a) and that 74.7 % (n=4225; 2418↑/1821↓) of differentially expressed genes were dysregulated in both, young and old compared to neonatal isolated cells (Fig. 2b,c,d). Accordingly, we could observe significant changes in old versus young mice (602↑/293↓) (Fig. 2e). Gene set enrichment analysis revealed differentially enriched hallmark pathways over the lifespan. Pro-inflammatory pathways including “TNFα signalling via NF-κB“ and interferon pathways significantly enriched in aged epithelial cells, highlighting an increased inflammatory signature with age (Fig. 2f). In parallel, pathways related to DNA damage repair, stemness and epithelial regeneration were decreased in adults in comparison with neonatal cells, consistent with the increase in DNA damage signalling observed in Fig 1. Furthermore, senescence-related gene sets were enriched in old versus neonatal cells (Suppl. Fig. 2a). Notably, WNT/β-catenin signalling was one of the hallmark gene sets that decreased over the lifespan. Expression patterns of selected genes related to regeneration, injury and aging highlighted a decrease in regeneration-associated markers (including *Sftpc*, *Hopx* and *Axin2),* an increased expression of injury associated transitional markers *(Lcn2, Sprr1a)* and an increased inflammatory (*Cxcl1, Lcn2* or *Csf2*) and ageing related (*Cdkn1a, Cdkn2a, Gdf15*) expression pattern in epithelial cells with increasing age (Fig.2g). In conclusion, these data demonstrate a decrease in regeneration and stemness pathways over lifespan and an increase in inflammatory and senescence-associated gene expression.

**Figure 2:**
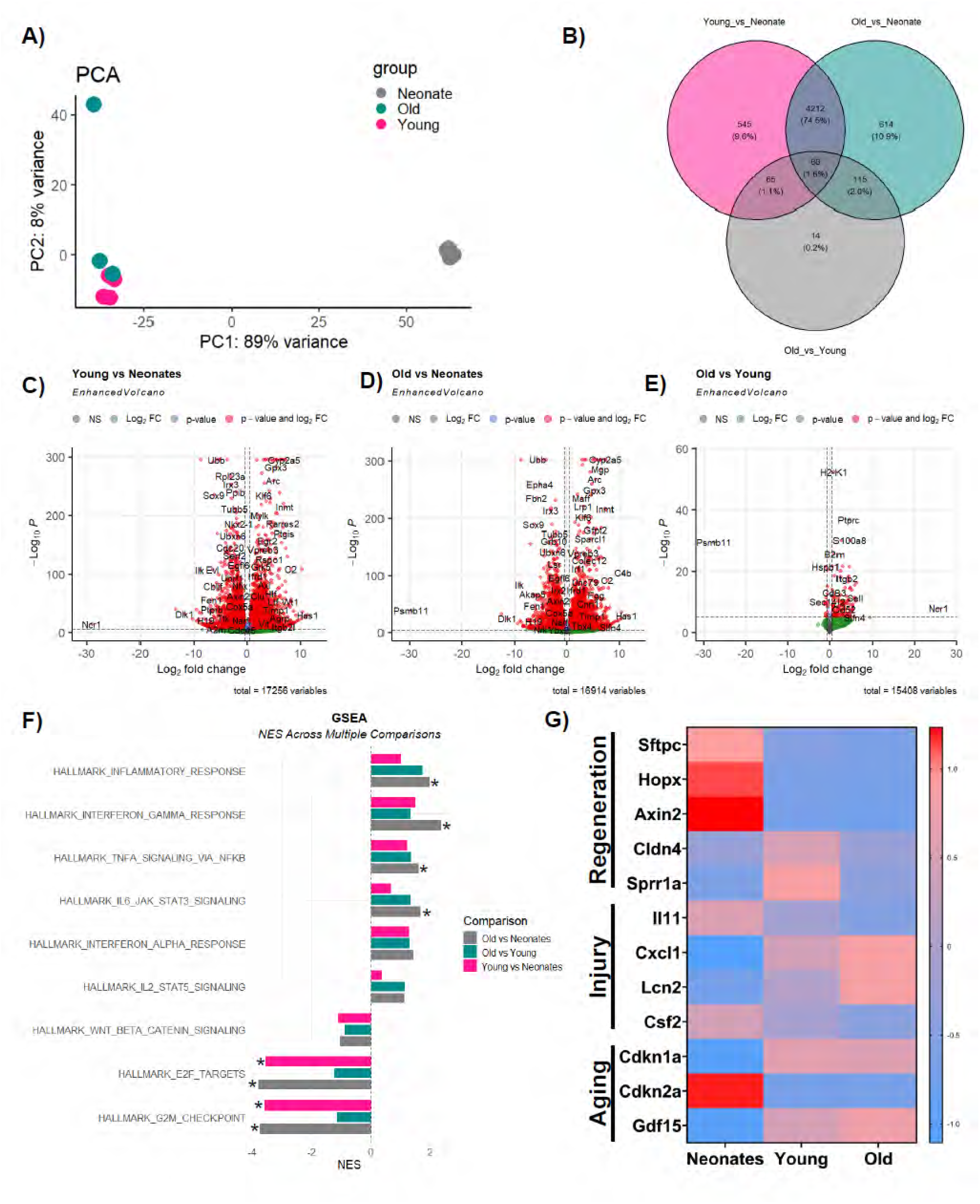
Neonatal Epithelium is intrinsically different from adults: **A**) Principal component analysis (PCA) of transcriptomes from primary lung EpCAM^+^ cells of neonatal, young and old mice shows age-dependent clustering. **B)** Venn Diagram showing overlap of differentially expressed genes between neonatal, young and old mice. **C, D, E)** Volcano plots visualizing differential gene expression between young and neonatal **(C)**, old and neonatal **(D)** and old and young **(E)** epithelial cells. Plots display log_2_ fold change against the adjusted P value (log_10_). **F)** Gene set enrichment analysis of top upregulated and downregulated pathways comparing young and neonatal epithelial cells (pink), old and neonatal epithelial cells (grey) and old and young epithelial cells(green). **G)** Heatmap of gene expression related to regeneration of the alveolar epithelium, injury response and aging compared between neonatal, young and old mouse lung epithelial cells. Statistical tests: **F)** fgsea in-built permutation-based statistical test. *\* p-value < 0.05*.

### Regenerative capacity of epithelial organoids decreases with age

Given the profound transcriptomic changes with age, we sought to test the functional regenerative capacity of lung epithelial cells over the lifespan using lung organoids. Neonatal, young, and old epithelial cells were used to establish organoid cultures that expressed markers of AT2 cells (proSP-C, Lamp3), alveolar differentiation intermediates (ADI; Krt8), and alveolar type I (AT1, Aqp5) cells (Fig. 3a), indicating the formation of alveolar organoids. In parallel, a subset of organoids stained positive for acetylated tubulin, suggesting the presence of airway-like organoids within the same cultures in neonatal and adult organoid cultures, consistent with previous reports using adult mouse EpCAM⁺ cells (Wu *et al)*.

**Figure 3:**
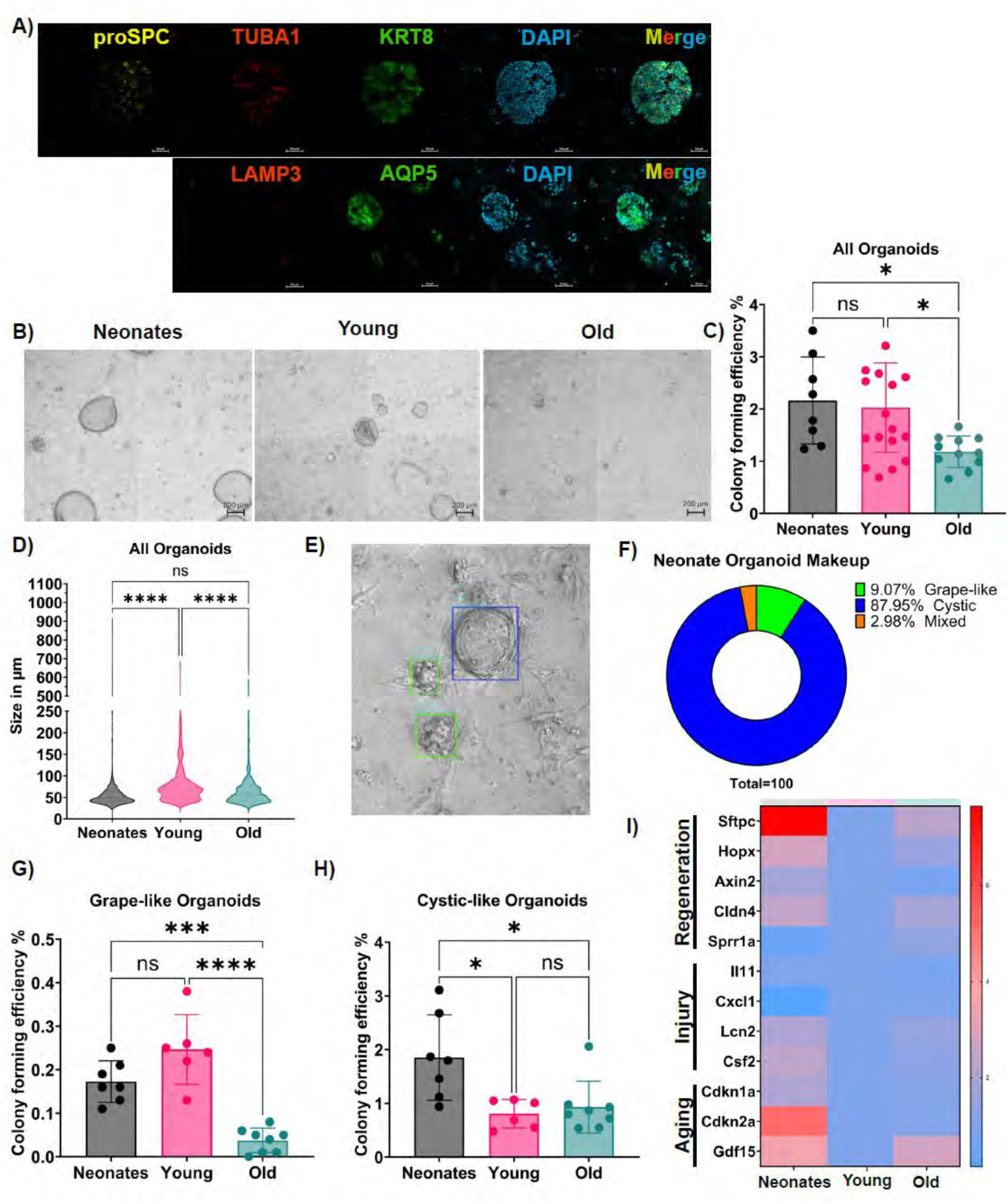
Organoids from neonates show differences compared to those from adults: **A**) Representative immunofluorescence images of neonatal lung organoids stained for pro-Surfactant Protein C (*pro-SPC*), Tubulin Alpha 1a (*Tuba1*), Cytokeratin 8 (*Krt8*), Lysosome-associated membrane glycoprotein 3 (*Lamp3*), Aquaporin 5 (*Aqp5*) and DAPI for nuclear staining. Scale bar: 50µm. **B)** Representative brightfield microscopy images of lung organoids from neonatal, young and old mice on Day 14 of culture. Scale bar: 200µm. **C, D)** Comparison of colony forming efficiency (%) and size (µm) of all lung organoids from neonatal, young and old mice. **E)** Representative image of the output from the Napari organoid counter showing the morphology of cystic (blue box) and grape (green box) like organoids. **F)** Pie chart showing the phenotypic makeup of lung organoids from neonates as annotated using the Napari organoid counter. **G, H)** Comparison of colony forming efficiency (%) and size (µm) of cystic-like and grape-like lung organoids from neonatal, young and old mice. **I)** Heatmap showing the expression of genes related to regeneration of the alveolar epithelium, injury response and aging analysed by RT-qPCR of lung organoids from neonatal, young and old mice. Statistical tests: **D, E, H, I)** Ordinary one-way ANOVA followed by Tukey’s multiple comparison test, *\* p-value < 0.05, ** p-value < 0.01, *** p-value < 0.001, **** p-value <0.0001*.

Age-dependent changes in organoid growth and morphology were quantified using high-content imaging combined with the Napari organoid counter (Bukas *et al*, 2024; Kastlmeier *et al*, 2023). Neonatal organoids exhibited a significantly higher colony-forming efficiency (CFE) compared to organoids derived from adult mice (Fig. 3b and c). In addition, organoid size varied across the lifespan (Fig. 3d). Given the coexistence of distinct organoid phenotypes across all age groups, organoids were further classified into grape-like (alveolar-like), cystic (airway-like), and mixed morphologies using the Napari organoid counter (Fig. 3e). Analysis of organoid composition revealed that neonatal cultures were predominantly composed of cystic organoids (Fig. 3f, Supplementary Figure 3f, g). With aging, grape-like organoids declined primarily from young to old mice (Fig. 3g), whereas cystic organoids already decreased from neonatal to young adult stages (Fig. 3h), suggesting differential age-related effects on distinct epithelial progenitor populations, as previously described (Rowbotham *et al*, 2023).

Gene expression analysis of cultured organoids largely recapitulated patterns observed in freshly isolated epithelial cells. Markers associated with alveolar epithelial regeneration, including *Sftpc*, *Hopx, Axin2*, and *Cldn4*, were reduced in adults compared to neonatal organoids (Fig. 3i), consistent with an overall decline in regenerative capacity (Lehmann *et al*, 2020). Notably, genes related to injury and aging were expressed at lower levels in young compared to both neonatal and old organoids. Interestingly, *Cdkn1a* (p21), a marker associated with senescence, was highly expressed in neonatal organoids compared to adult organoids. While most findings were consistent with gene expression in freshly isolated epithelial cells, senescence-associated signatures appeared less pronounced in organoid cultures, potentially due to competitive outgrowth of non-senescent cells during *in vitro* expansion (Fig. 3i).

Finally, the pro-inflammatory epithelial cytokine *Cxcl1* was more highly expressed in organoids derived from adult mice compared to neonatal counterparts, suggesting that age-associated inflammatory features are at least partially retained in epithelial organoid cultures.

### Inflammaged niche controls regenerative capacity

We next investigated whether the aged epithelial niche contributes to the observed decline in regenerative capacity. To this end, young organoids were cultured in conditioned medium derived from either young or aged organoids (Fig. 4a). Notably, exposure to conditioned medium from aged organoids significantly reduced CFE of young epithelial cells (Fig. 4b). This effect was most pronounced in alveolar, grape-like organoids, while the fraction of cystic organoids remained unchanged (Fig. 4c, d). This suggests that a paracrine mechanism mediated by a secreted factor can influence regenerative capacity primarily in the alveolar cells.

**Figure 4:**
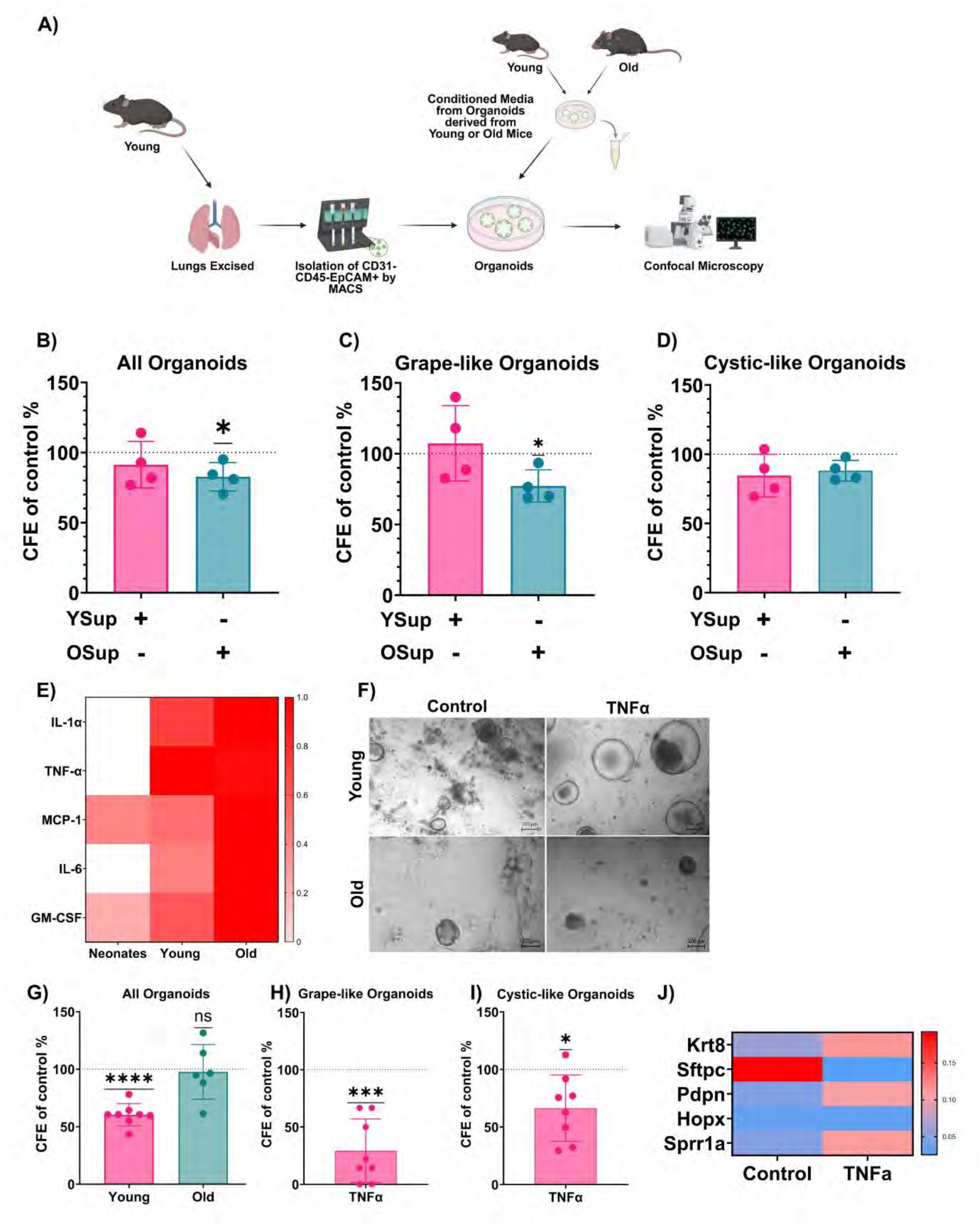
Inflammatory stimuli influence organoid formation across age groups: **A**) Scheme explaining the experimental workflow of lung organoids from young mice treated with conditioned media from organoids from young and old mice. **B, C, D)** Comparison of colony forming efficiency as a percentage of untreated control of all, grape-like and cystic-like lung organoids from young mice treated with conditioned media. **E)** Heatmap showing the levels of released pro-inflammatory cytokines as measured by LegendPlex Multiplex ImmunoAssay of supernatants of lung organoids from neonatal, young and old mice. **F)** Representative brightfield microscopy images of lung organoids young and old mice treated with TNF-a on day 14 of culture. **G)** Comparison of colony forming efficiency as a percentage of untreated control of all lung organoids from young and old mice treated with TNF-a. **H, I)** Comparison of colony forming efficiency as a percentage of control of grape-like and cystic-like lung organoids from young mice treated with TNF-a. **J)** Heatmap showing expression of genes related to regeneration of the alveolar epithelium analysed by RT-qPCR of lung organoids from young mice treated with TNF-a. Statistical tests: **B, C, D, G, H, I)** One sample t-test, *\* p-value < 0.05, ** p-value < 0.01, *** p-value < 0.001, **** p-value <0.0001*.

Among secreted factors, cytokines are key regulators of epithelial cell function during homeostasis and injury, and epithelial cells themselves represent an important source of pro-inflammatory mediators, particularly in the context of aging. Accordingly, the gene expression of various age-related senescence-associated secretory phenotype (SASP) factors were upregulated in epithelial cells over the lifespan (Suppl. Fig. 2b). We therefore assessed whether cytokine secretion is altered in organoid cultures across the lifespan. Profiling of culture supernatants revealed detectable levels of MCP-1 and GM-CSF in all organoid conditions (Fig. 4e). Importantly, several inflammatory cytokines, including IL-1α, TNF-α, MCP-1, IL-6, and GM-CSF, were significantly increased in supernatants from organoids derived from aged epithelial cells, suggesting that these cultures recapitulate features of an inflammaged niche (Fig. 4e).

Given that TNF-α was strongly upregulated with age and has been described as a potent regulator of stem cell function, we next examined its direct effect on epithelial regenerative capacity (Fig. 4f). In young organoid cultures, exogenous TNF-α significantly reduced CFE (Fig. 4g). In contrast, no additional effect was observed in organoids derived from aged epithelial cells, likely due to already elevated endogenous TNF-α levels. In young cultures, TNF-α treatment predominantly affected grape-like organoids (Fig. 4h, i), further supporting a selective effect on alveolar epithelial cells. This was accompanied by a shift in cellular identity, with reduced AT2 and increased ADI gene expression in treated organoids (Fig. 4j), consistent with impaired alveolar regeneration.

### Disease-relevant stimuli influence organoid formation

To assess age-dependent susceptibility to disease-relevant stimuli, organoids across the lifespan were exposed to TGF-β and IL-1β, mediators that are induced in multiple chronic lung diseases including IPF and COPD. TGF-β1 markedly impaired organoid growth and reduced size in all age groups (Suppl. Fig. 4a, b, c), whereas IL-1β increased organoid formation across ages, with a significant size increase only in aged organoids (Suppl. Fig. 4a, d, e). At the transcriptional level, TGF-β1 downregulated regeneration– and injury-associated genes (e.g., *Krt8, Lcn2*) while inducing WNT target genes (*Wisp1, Axin2*), particularly in neonatal organoids (Suppl. Fig. 4f). In contrast, IL-1β had limited effects on WNT signalling but induced injury-related genes, with a stronger response in neonatal organoids. Overall, while organoid formation responded similarly across ages, neonatal organoids showed greater transcriptional plasticity and aged organoids exhibited enhanced pro-inflammatory responses.

### Decline in regenerative capacity over the lifespan is epigenetically imprinted

To determine which age-associated transcriptional changes are maintained in organoid cultures, we performed bulk RNA sequencing of organoids derived from young and old mice (Fig. 5a) and compared these profiles to those of freshly isolated epithelial cells (data from Fig. 2). Aged organoids exhibited a substantial number of differentially expressed genes compared to young organoids after 14 days in culture (Fig. 5b).

**Figure 5:**
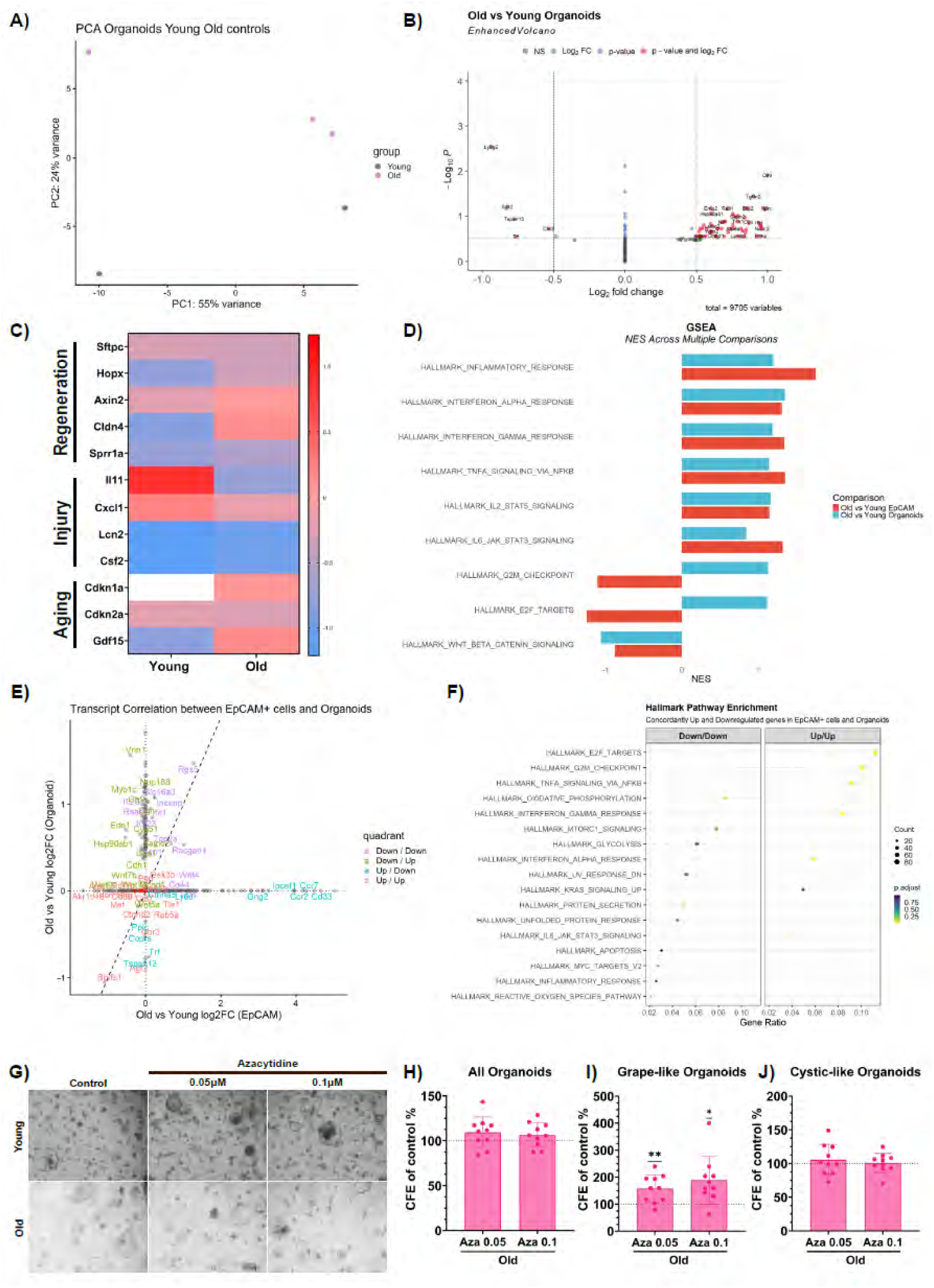
Transcriptional and translational changes in organoids across the lifespan upon injury: **A**) Principal Component Analysis of Bulk Sequencing of Organoids from young and old mice. **B)** Volcano plots visualizing differential gene expression between organoids from young and old mice. Plots display log_2_ fold change against the adjusted P value (log_10_). **C)** Heatmap of gene expression related to regeneration of the alveolar epithelium, injury response and aging compared between young and old mouse lung organoids as measured by Bulk RNA sequencing. **D)** Gene set enrichment analysis of top upregulated and downregulated pathways comparing old vs. young EpCAM^+^ cells (dark orange) and old vs. young organoids (cyan). **E)** Plot showing correlation of expression of selected genes in old vs young EpCAM^+^ vs old vs young Organoids. **F)** Gene Set Enrichment Analysis of genes concordantly up and downregulated in EpCAM^+^ cells and organoids from young and old mice. **G)** Representative brightfield microscopy images of lung organoids from young and old mice treated with Azacytidine on Day 14 of culture. **H, I, J)** Comparison of colony forming efficiency (%) of all grape-like and cystic-like lung organoids from old mice treated with Azacytidine. Statistical tests: **H, I, J)** One sample t-test, *\* p-value < 0.05, ** p-value < 0.01, *** p-value < 0.001, **** p-value <0.0001*.

Focusing on key regenerative and injury-associated genes, we observed sustained upregulation of injury markers (*Cldn4, Sprr1a, Lcn2*) and aging-associated genes (*Cdkn1a, Gdf15*), alongside persistent downregulation of the AT2 marker *Sftpc* in aged organoids (Fig. 5c). Gene set enrichment analysis further demonstrated that similar pathways are regulated upon aging in both freshly isolated epithelial cells and organoid cultures (Fig. 5d). In particular, inflammatory pathways were upregulated, whereas pathways related to DNA repair, cell cycle regulation, and developmental signalling, including WNT/β-catenin signalling, were downregulated.

To identify age-related gene programs that are preserved during organoid culture in an unbiased manner, we compared differentially expressed genes between young and old EpCAM⁺ epithelial cells and their corresponding organoids, focusing on genes regulated in the same direction (up/up or down/down; Fig 5e). This analysis revealed enrichment of inflammatory pathways, including NF-κB signalling, among consistently upregulated pathways (Fig 5f). Conversely, multiple metabolic pathways were downregulated in both epithelial cells and organoids, indicating that key aspects of age-associated transcriptional dysregulation are retained in organoid cultures.

Given the persistence of these aging-associated signatures, we hypothesized that epigenetic mechanisms may contribute to their maintenance. DNA methylation, a stable epigenetic modification associated with gene silencing, has been implicated in aging-related processes (Kurbanov *et al*, 2025; Hernandez Cordero & Leung, 2024; Wang *et al*, 2022). To test whether DNA methylation contributes to the reduced regenerative capacity of aged organoids, we treated adult organoid cultures with the DNMT inhibitor 5-azacytidine (Fig 5g). This treatment resulted in a significant increase in colony-forming efficiency in grape-like organoids, suggesting that increased DNA methylation contributes to impaired alveolar regeneration with age (Fig 5h-j).

Notably, although regenerative capacity was partially restored, the expression of inflammatory mediators remained largely unchanged following 5-azacytidine treatment (Suppl. Fig. 4g) indicating a potential epigenetic uncoupling between inflammatory signalling and regenerative decline in aged epithelial cells.

### Activating WNT/ β-catenin rescues regenerative capacity

Neonatal organoids exhibited enhanced regenerative capacity, correlating with increased WNT activity in both freshly isolated epithelial cells and organoid cultures (Fig. 5d). In freshly isolated cells, WNT ligands as well as bona fide target genes were highly expressed in neonates, whereas their expression was downregulated upon aging (Fig. 6a). Here, molecular WNT signatures between young and old adult cells were comparable. To determine whether WNT activation could restore regenerative capacity in adult organoids to mimic the neonatal state, cultures were treated with LiCl, a GSK3β inhibitor. LiCl treatment significantly increased *Axin2* expression in both young and old organoids, confirming activation of WNT/β-catenin signalling (Fig. 6b).

**Figure 6:**
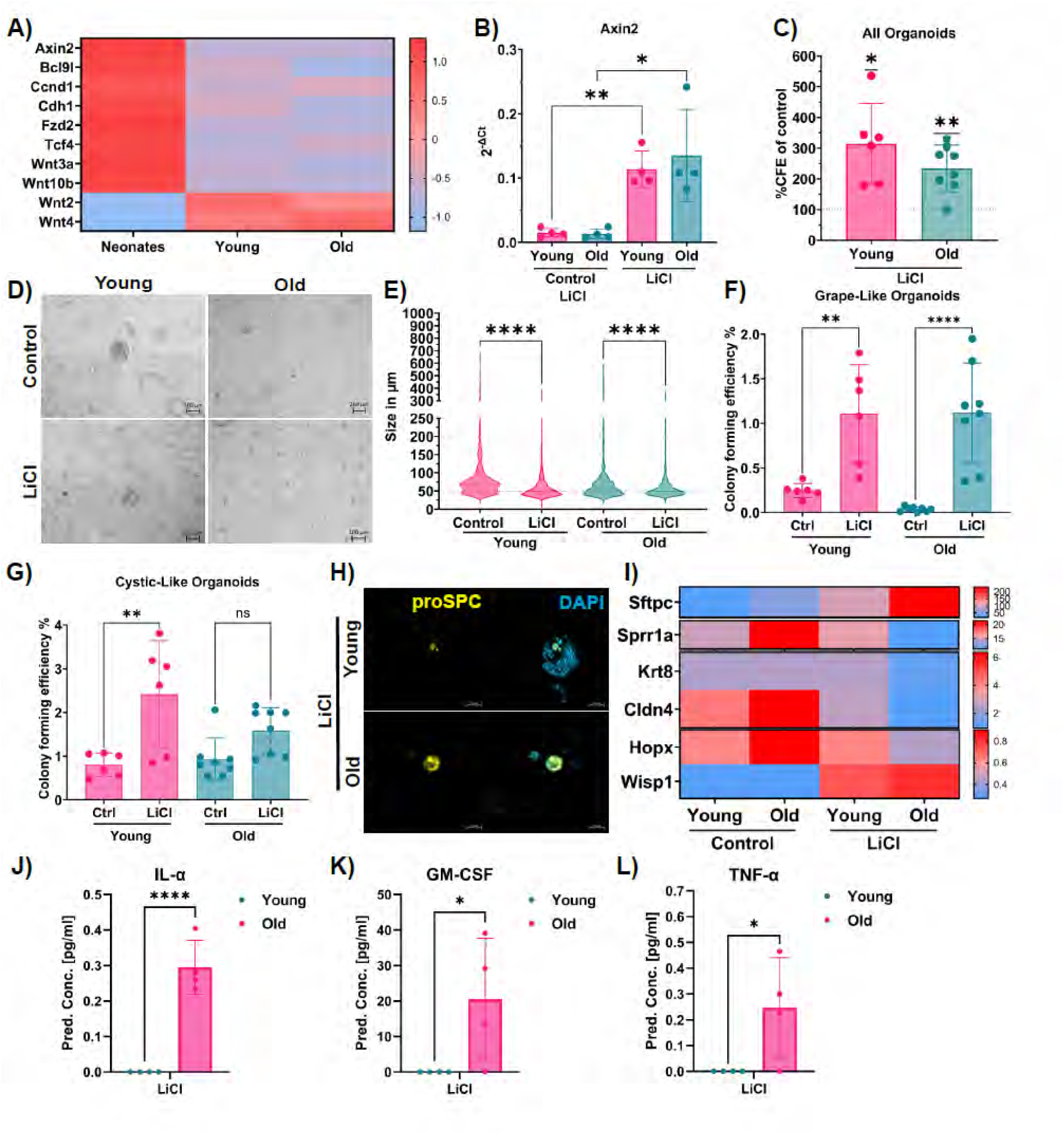
Regenerative capacity of epithelial organoids decreases with age and can be restored by activating Wnt signalling: **A**) Heatmap showing the expression of genes related to Wnt signalling in EpCAM+ cells from neonatal, young and old mice as measured using Bulk RNA sequencing. **B)** Relative gene expression of Axin2 to Hprt as analysed by RT-qPCR in untreated and LiCl treated organoids grown for 14 days from young and old mice. **C)** Comparison of colony forming efficiency (%) of all lung organoids from young and old mice treated with LiCl. **D)** Representative brightfield microscopy images of control and LiCl treated lung organoids from young and old mice on day 14 of culture. Scale bar: 200 µm **E)** Comparison of organoid size (µm) of all control and LiCl treated lung organoids from young and old mice. **F, G)** Comparison of colony forming efficiency (%) of grape-like and cystic-like control and LiCl treated lung organoids from young and old mice**. H)** Representative Immunofluorescence images of LiCl treated lung organoids from young and old mice stained for pro-Surfactant Protein C (pro-SPC) (yellow) and DAPI (blue) for nuclear staining. Scale bar: 50µm. **I)** Heatmap showing expression of genes related to regeneration of the alveolar epithelium and Wnt signalling analysed by RT-qPCR of untreated and LiCl treated lung organoids from young and mice. **J, K, L)** Bar plots showing the levels of IL-1α, GM-CSF and TNF-α in the supernatants of LiCl treated organoids from young and old mice as measured by LegendPlex Multiplex ImmunoAssay. Statistical tests: **B)** Unpaired t-test with Welch’s correction. *\* p-value < 0.05, ** p-value < 0.01, *** p-value < 0.001, **** p-value <0.0001.* **C, D, F, G, H, K, L)** Ordinary one-way ANOVA followed by Tukey’s multiple comparison test. *\* p-value < 0.05, ** p-value < 0.01, *** p-value < 0.001, **** p-value <0.0001*.

Functionally, WNT activation increased CFE in both age groups while reducing organoid size (Fig. 6c, d, e). This effect was most pronounced in grape-like organoids, indicating a preferential impact on alveolar structures (Fig. 6f). Consistently, LiCl treatment enhanced pro-SPC expression in both young and old organoids (Fig. 6h), supporting an effect on alveolar progenitors. LiCl treatment decreased markers of differentiation intermediates, suggesting a reprogramming towards a more stem-like state (Fig. 6i).

Notably, despite the observed WNT-associated pro-stemness effects, pro-inflammatory cytokine levels remained elevated in organoids derived from aged mice (Fig. 6j,k,l, Suppl. Fig. 5c,d,e), suggesting that inflammatory signalling is not regulated by WNT/β-catenin and further supporting an uncoupling of impaired regeneration and inflammaging.

## Discussion

Here we show that lung epithelial cells exhibit differences in regeneration, gene expression, and cytokine production across the lifespan. Organoid cultures recapitulate these features and enable dissection of stem cell–intrinsic and inflammatory niche effects. Our data suggest that inflammaging and epigenetic imprinting both contribute to age-related regenerative decline and can be experimentally uncoupled.

Using an organoid system derived from lung epithelial progenitors, containing a mix of bronchiolar and alveolar stem cells as described previously (Wu *et al)*, combined with a machine-learning-assisted morphological counting tool, we decoupled age-related CFE effects on each compartment separately (Bukas *et al*, 2024). Using this tool, we are the first to compare neonatal lung epithelium with young and old epithelium. Consistent with previous studies, regenerative capacity of the alveolar cells declines between young and late adulthood (Strunz *et al*, 2020; Man *et al*, 2026; Rowbotham *et al*, 2023); however, there is a negligible decline between neonatal and young alveolar organoids. While prior work describes a preferential regenerative decline in alveolar over bronchiolar progenitors (Hopkinson *et al*, 2024; Duan *et al*, 2021; Kapellos *et al*, 2023), we show here, that bronchiolar epithelium undergoes a sharper decline from neonatal stages into adulthood. This suggests a differential regulation of regenerative capacity of specific epithelial stem cells over the lifespan. Epigenetic mechanisms may contribute to the regenerative decline with age (Rowbotham *et al*, 2023; Baker *et al*, 2026). Treatment with Azacytidine, an inhibitor of de novo DNA methyltransferases increased alveolar regenerative capacity, suggesting that DNA methylation controls loss of regenerative capacity over the lifespan and can be reversed by inhibiting *de novo* methylation. Studying the effects of other epigenetic marks including histone modifications on driving regenerative decline would be of great interest (Rowbotham *et al*, 2023).

Immune cells link inflammation to regenerative decline, with inflammaging-associated cytokines such as GZMK and IFN-γ shown to drive this decline in lung epithelium (Kapellos *et al*, 2023; Dost *et al*, 2026; Jensen *et al*, 2026; Gote-Schniering *et al*, 2025; Rustam *et al*, 2023; Glisinski *et al*, 2020). Here, we show that organoids from aged mice establish an inflammaged niche, suggesting that part of the (local) inflammaged milieu can be driven by epithelial cell aging (Kathiriya *et al*, 2022; Toscano-Marquez *et al*, 2026). This niche influences regenerative capacity since culturing young epithelial cells in an inflammaged niche of old organoids reduced their regenerative capacity. Similarly, the presence of TNF-α, one of the main inflammation-associated cytokines in the organoid cultures, reduced regenerative capacity, highlighting the susceptibility of epithelial cells to their microenvironment. Notably, inhibition of DNMTs did not affect inflammation, suggesting that reduced regenerative decline and epithelial inflammaging can be decoupled as aging hallmarks experimentally and epigenetic reprogramming can overcome the detrimental effect of an inflammaged niche. How inflammatory imprinting that is maintained in the organoid cultures is epigenetically maintained *in vitro* independently of DNA methylation warrants further investigation.

Using epithelial lung organoids allowed us to focus on the role of the epithelium and epithelial-driven inflammation over the lifespan in vitro. At the same time, this approach did not allow us to study native cell-cell interactions and mechanics, including immune responses or remodeling at the tissue level. Supplementing immune cells in the organoid setup or the use of more complex models such as precision-cut lung slices could be a next step to study these effects in a tissue context (Lehmann *et al*, 2025).

Over the lifespan, the largest differences found in our study occur between neonatal and adult cells, indicating rapid early-life reprogramming of the lung epithelium. Consistent with recent human studies, aging may involve discrete transition points marked by abrupt functional decline rather than gradual deterioration (Lehallier *et al*, 2019; Shen *et al*, 2024). Our analysis of three timepoints, together with prior studies, captures age-associated changes in the lung epithelium, revealing distinct functional and molecular states and marked early-life alterations (Baker *et al*, 2026; Han *et al*, 2023; Melo-Narváez *et al*, 2020; Man *et al*, 2026). Longitudinal studies are needed to determine whether these changes are progressive or reflect discrete transition phases.

Here we focused on activating WNT/β-catenin in the adult lung organoids, a developmental pathway that was highly enriched in the neonatal cells but downregulated in the adult cells. The WNT signalling pathway is a central regulator of lung development and regeneration (Nabhan *et al*, 2018; Frank *et al*, 2016; Lee *et al*, 2017; Hu *et al*, 2021), yet its role in aging-associated lung disease appears complex and context-dependent (Kneidinger *et al*, 2011; Nabhan *et al*, 2023). Here, we observe a reduction in canonical WNT/β-catenin in aged lung epithelial cells, a phenotype that is retained in organoid cultures, suggesting a stable, potentially epigenetically imprinted suppression of WNT signalling. Notably, reactivation of WNT signalling in organoids derived from aged epithelial cells enhanced regenerative capacity, particularly by promoting alveolar epithelial organoid formation. This supports recent studies demonstrating beneficial effects of WNT activation in chronic lung disease (Nabhan *et al*, 2023; Costa *et al*, 2021). However, these findings must be interpreted in light of the dual role of WNT signalling in aging tissues (Lehmann *et al*, 2016). Sustained or excessive activation of WNT/β-catenin has been linked to the induction of cellular senescence, and we have identified a subset of WNT-high cells in the aged lung, indicating that precise regulation of this pathway activity is critical (Lehmann *et al*, 2020). In our model, WNT activation via GSK3β inhibition using lithium chloride improved regenerative output but did not attenuate inflammatory cytokine release, suggesting that epithelial regeneration and inflammatory signalling can be uncoupled. Given that GSK3β regulates multiple signalling pathways (Beurel *et al*, 2015), these findings further highlight the need for more targeted approaches to modulate WNT signalling. Intriguingly, retrospective clinical observations suggesting improved emphysematous phenotypes in lithium-treated COPD patients (Chandra *et al*, 2025) raise the possibility that modulation of WNT signalling may enhance epithelial stem cell function *in vivo*. Additionally, lithium may have anti-aging effects, given that its use has been associated with a reduction of aging biomarkers including telomere length (Courtes *et al*, 2024). Together, these data support a model in which transient and carefully titrated activation of WNT signalling promotes epithelial regeneration, whereas sustained (over)activation may contribute to senescence and maladaptive remodeling as observed in pulmonary fibrosis (Königshoff *et al*, 2008; Patel *et al*, 2024).

Together, our findings demonstrate that lung epithelial cells undergo profound, early changes across the lifespan, with regenerative decline driven by both intrinsic epigenetic imprinting and epithelial-derived inflammaging that can be experimentally uncoupled in vitro. These insights highlight the need for combinatorial therapeutic strategies that simultaneously restore epithelial plasticity and modulate inflammatory signalling to promote effective lung regeneration during aging.

## Author contribution

ML and CV conceptualized the study. GB, EMG, CS, MCMN, CB, TN, AM, FFB, LMT, QH, HSE and EFP designed and contributed to the design and conducted experiments. MP and FC developed the Napari Organoid Counter. GB and MCMN performed bioinformatic analyses. RMW and ALJ contributed to conceptualization through fruitful scientific discussions that shaped the experimental approach and improved the manuscript. AH, MP, BS, MK, TS, MAAA, AÖY, CV and ML provided funding. GB, EMG, CV and ML wrote the first manuscript draft. GB and ML wrote the final manuscript version. All authors have read and approved the final manuscript.

## Acknowledgments

We would like to express our sincere gratitude to Kathrin Federl and Max Mikado Götz for excellent technical assistance regarding primary cell isolation and cultures. We would also like to acknowledge Eva Herker for support with imaging organoid cultures. She acknowledges support from the Deutsche Forschungsgemeinschaft – 517270163. ML acknowledges support from the Deutsche Forschungsgemeinschaft (DFG, German Research Foundation) – 512453064 and 542102810 and the von Behring Röntgen Foundation (71_0011), and the Hessisches Ministerium für Wissenschaft und Forschung, Kunst und Kultur (LOEWE Habitat). CV acknowledges support from the Deutsche Forschungsgemeinschaft (DFG, German Research Foundation) – 530370007. CV, TS, ML, BS report funding from the German Center for Lung Research (DZL).

## Supplementary Figures

**Supplementary Figure 1:**
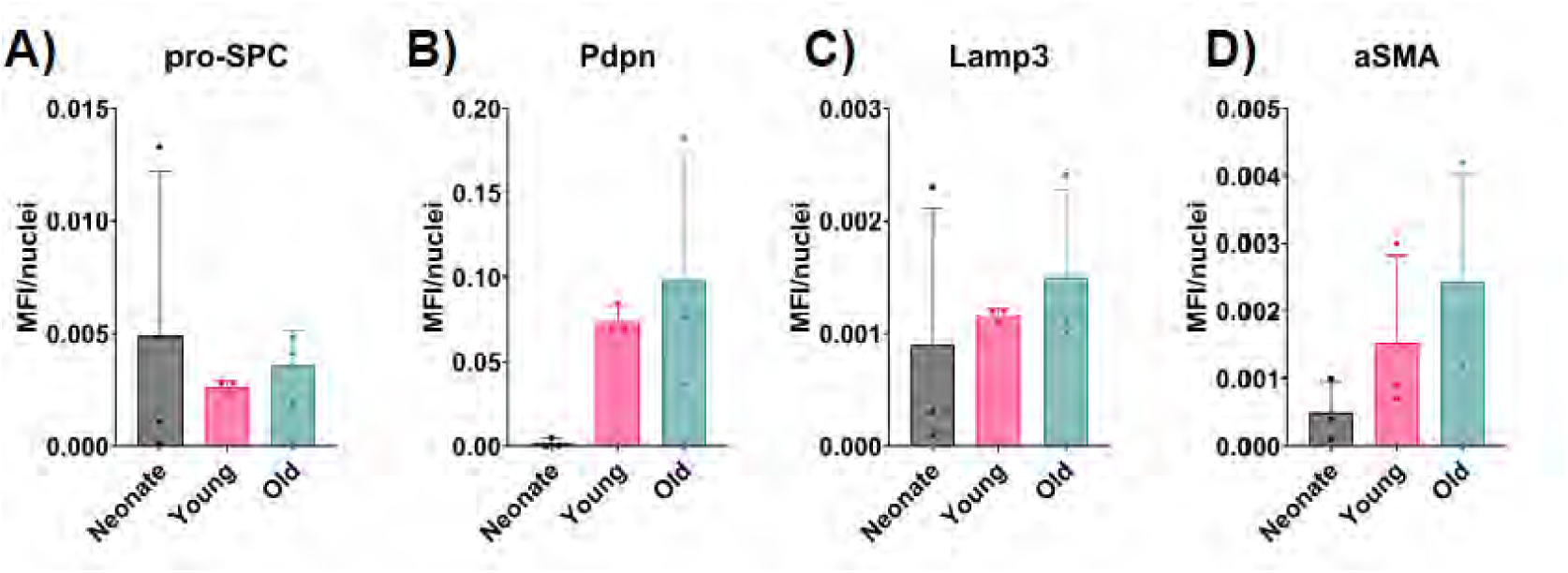
**A, B, C, D**) Quantification of mean fluorescence intensity normalized to nuclei of pro-SPC, PDPN, LAMP3 and aSMA in FFPE sections of lungs from neonatal, young and old mice. Statistical tests: **A, B, C, D)** Ordinary one-way ANOVA followed by Tukey’s multiple comparison test. No significance detected.

**Supplementary Figure 2:**
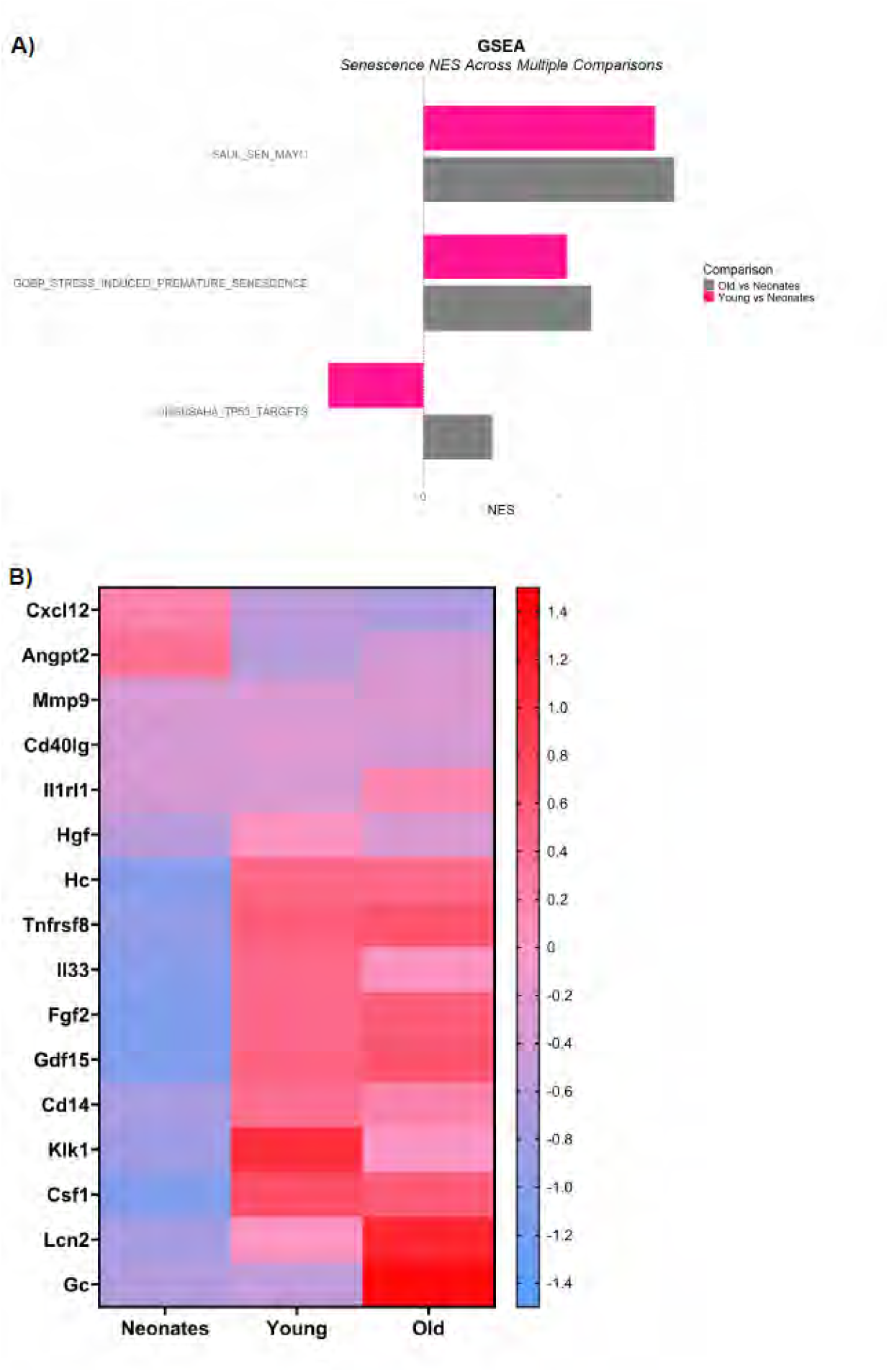
**A**) Gene set enrichment analysis of top upregulated and downregulated gene sets related to senescence comparing young and neonatal epithelial cells (pink) and old and neonatal epithelial cells (grey). **B)** Heatmap showing the expression of genes related to SASP in EpCAM+ cells from neonatal, young and old mice.

**Supplementary Figure 3:**
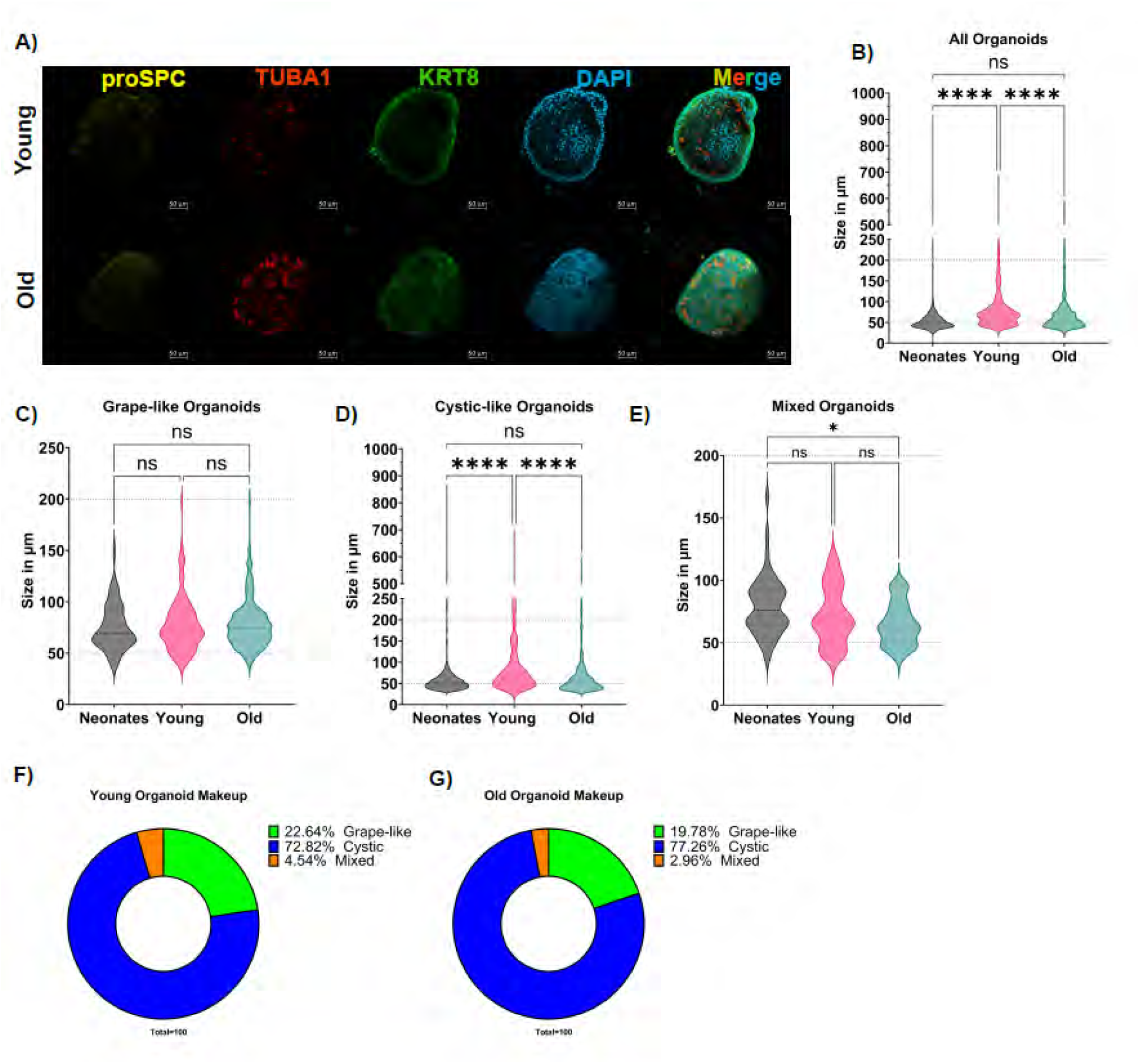
**A**) Representative images of immunofluorescence staining of mouse lungs from young and old mice stained for pro-SPC (yellow), TUBA1A (red), KRT8 (green) and DAPI (blue). Scale bar = 50 µm. **B)** Comparison of organoid Size (µm) of all lung organoids from neonatal, young and old mice. **C, D, E)** Comparison of organoid Size (µm) of grape-like, cystic-like and mixed lung organoids from neonatal, young and old mice. **F, G)** Pie chart showing the phenotypic makeup of lung organoids from young and old mice as annotated using ML-assisted Organoid Counting. Statistical tests: **B, D, E)** Ordinary one-way ANOVA followed by Tukey’s multiple comparison test. *\* p-value < 0.05, ** p-value < 0.01, *** p-value < 0.001, **** p-value <0.0001.* **C)** Ordinary one-way ANOVA followed by Holm-Sidak’s multiple comparison test. No significance detected.

**Supplementary Figure 4:**
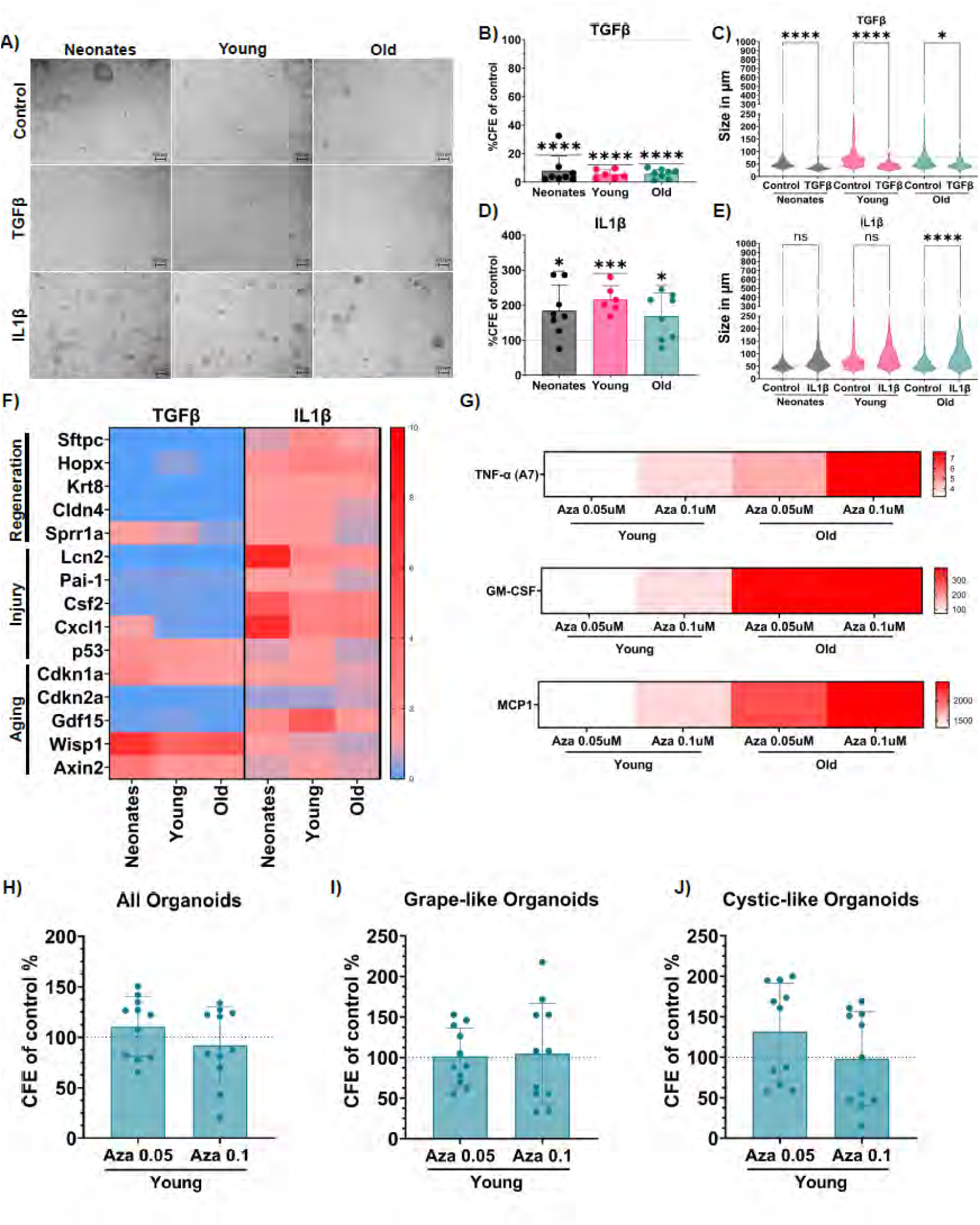
**A**) Representative brightfield images of control, TGF-β1 and IL1β treated lung organoids from neonatal, young and old mice. **B, D)** Comparison of colony forming efficiency of control, TGF-β1 and IL1β treated lung organoids as a percentage of control from neonates, young and old mice. **C, E)** Comparison of organoid size (µm) of control, TGF-β1 and IL1β treated lung organoids from neonates, young and old mice. **F)** Heatmap showing the expression of genes related to regeneration of the alveolar epithelium, injury response and aging analysed by RT-qPCR of control, TGF-β1 and IL1β treated lung organoids from neonatal, young and old mice. **G)** Heatmap showing the levels of released pro-inflammatory cytokines as measured by LegendPlex Multiplex ImmunoAssay in supernatants of lung organoids treated with Azacytidine from young and old mice. **H, I, J)** Comparison of colony forming efficiency as a percentage of control of all, grape-like and cystic-like lung organoids from young mice treated with Azacytidine. Statistical tests: **C, E)** Ordinary one-way ANOVA followed by Sidak’s multiple comparison test, *\* p-value < 0.05, ** p-value < 0.01, *** p-value < 0.001, **** p-value <0.0001.* **B, D, H, I, J)** One sample t-test, *\* p-value < 0.05, ** p-value < 0.01, *** p-value < 0.001, **** p-value <0.0001*.

**Supplementary Figure 5:**
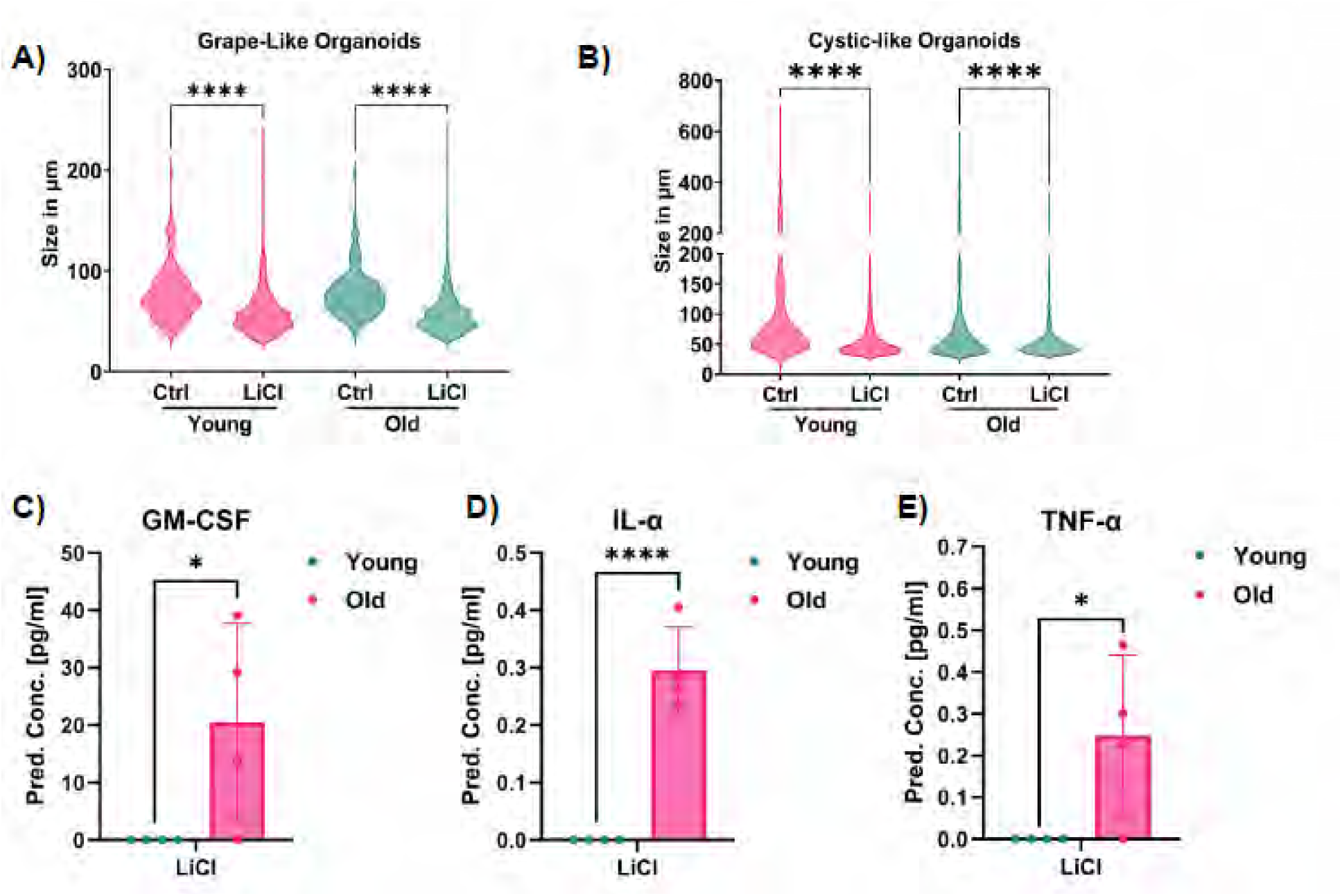
**A, B**) Comparison of organoid size (µm) of grape-like and cystic-like control and LiCl treated lung organoids from young and old mice. **C, D, E)** Bar Plots showing the levels of released pro-inflammatory cytokines as measured by LegendPlex Multiplex ImmunoAssay in supernatants of lung organoids treated with LiCl from young and old mice. Statistical tests: **A, B, C, D, E)** Ordinary one-way ANOVA followed by Tukey’s multiple comparison test. *\* p-value < 0.05, ** p-value < 0.01, *** p-value < 0.001, **** p-value <0.0001*.

